# *Vibrio parahaemolyticus* T6SS2 effector repertoires

**DOI:** 10.1101/2022.11.17.516722

**Authors:** Daniel Tchelet, Kinga Keppel, Eran Bosis, Dor Salomon

**Author notes:** Address correspondence to Dor Salomon.

## Abstract

All strains of the marine bacterium *Vibrio parahaemolyticus* harbor a type VI secretion system (T6SS) named T6SS2, suggesting that this system plays an important role in the life cycle of this emerging pathogen. Although T6SS2 was recently shown to play a role in interbacterial competition, its effector repertoire remains unknown. Here, we employed proteomics to investigate the T6SS2 secretome of two *V. parahaemolyticus* strains, and we identified several antibacterial effectors encoded outside of the main T6SS2 gene cluster. We revealed two T6SS2-secreted proteins that are conserved in this species, indicating that they constitute the core secretome of T6SS2; other identified effectors are found only in subsets of strains, suggesting that they comprise an accessory effector arsenal of T6SS2. Remarkably, a conserved Rhs repeat-containing effector serves as a quality control checkpoint and is required for T6SS2 activity. Our results reveal the effector repertoire of a conserved T6SS, some of which have no known activity and have not been previously associated with T6SSs.

## Introduction

Members of the *Vibrionaceae* family are aquatic, Gram-negative bacteria (1). Many *Vibrio* species are pathogens of humans and marine animals (2–4). New pathogenic strains emerge due to the natural competency of these bacteria and due to horizontal gene transfer, enabling the acquisition of new virulence traits (5, 6).

In the marine environment, vibrios are in constant competition with rival bacteria over resources (7, 8). They also interact with protists that either prey on them or that serve as a replicative niche (9–11). Interactions with both bacteria and eukaryotes can be mediated by the type VI secretion system (T6SS), which is employed by many vibrios (12). T6SS is a molecular nanomachine that delivers toxic proteins, called effectors, into neighboring cells (13, 14). A missile-like structure, comprising an inner tube composed of stacked hexameric rings of Hcp proteins capped by a spike consisting of a VgrG protein trimer sharpened by a PAAR repeat-containing protein, is propelled out of the cell and into an adjacent cell by a contractile sheath that engulfs it (15, 16). The tube-spike complex is decorated with two types of effectors: (1) specialized effectors, which are the structural components Hcp, VgrG, or PAAR containing a C-terminal toxin domain extension; and (2) cargo effectors, which are proteins that non-covalently bind to a structural protein (17), either directly or aided by an adaptor protein (18), a tether (19), or a co-effector (20). Most T6SSs investigated to date deliver effectors with antibacterial activities and have therefore been implicated in interbacterial competitions (7, 21—24). However, several *Vibrio* T6SSs have also been shown to deliver effectors that target eukaryotic cells, indicating that they also mediate interactions with eukaryotes (13, 23, 25, 26). Notably, antibacterial effectors are encoded in bi-cistronic units together with cognate immunity proteins that antagonize their toxic activity to prevent self or kin-intoxication (14, 27).

*Vibrio parahaemolyticus* is a leading cause of gastroenteritis caused from consuming undercooked or raw seafood (3). It is also the major cause of acute hepatopancreatic necrosis disease (AHPND) in shrimp (28). The molecular determinant that governs *V. parahaemolyticus* virulence against humans has been identified as a type III secretion system (29, 30); the virulence determinant against shrimp was identified as the PirA/B toxin (31). *V. parahaemolyticus* also harbors T6SSs (32). T6SS1, which is found in most but not in all strains (32), mediates antibacterial toxicity during interbacterial competition by delivering effector repertoires that differ between strains (32–35). T6SS1 was investigated in several strains and was shown to be active under warm marine-like conditions (i.e., 3% [wt/vol] NaCl at 30°C) upon surface sensing activation (21). We recently showed that all *V. parahaemolyticus* strains harbor a conserved T6SS, named T6SS2, suggesting that this system plays an important role in the life cycle of this pathogen (32). Although recent reports indicated that T6SS2 mediates antibacterial activities (35, 36), its effector repertoire remains unknown.

In this work, we investigated T6SS2 in two *V. parahaemolyticus* strains: the reference clinical strain RIMD 2210633 (37) and the environmental strain BB22OP (38). We showed that T6SS2 in both strains plays a role in interbacterial competition. Using comparative proteomics, followed by experimental validations, we revealed the effector repertoire of T6SS2 in both strains. We found that *V. parahaemolyticus* T6SS2 employs a conserved Rhs repeat-containing effector that is essential for its activity, as well as accessory effectors that differ between strains.

## Results

### Identifying the T6SS2 secretome in *V. parahaemolyticus* BB22OP

We previously showed that T6SS2 in *V. parahaemolyticus* BB22OP plays a role in interbacterial competition under warm, marine-like conditions (i.e., 3% [wt/vol] NaCl at 30°C) (35), suggesting that this system delivers antibacterial effectors. To reveal the secretome of this T6SS2 and to identify its effectors, we used mass spectrometry analysis and compared the proteins secreted by a wild-type strain with those secreted by a strain in which we inactivated T6SS2 (T6SS2^-^; Δ*hcp2*). We identified eight proteins that were significantly enriched in the secretome of the wild-type (T6SS2^+^) strain (**Fig. 1A**, **Table 1,** and **Supplementary Dataset S1**). These proteins include the three secreted structural components of the T6SS tube-spike complex, Hcp2, VgrG2, and PAAR2 (16), as well as five additional proteins. These additional proteins are predicted to be antibacterial effectors, since they are encoded next to predicted cognate immunity proteins (**Fig. 1B**): Two have a predicted Lipase_3 domain (WP_015313641.1 / T2LipA^BB22^ and WP_015296300.1 / T2LipB^BB22^; they are 36.6% identical across the entire sequence), one has a membrane-disrupting Tme domain (WP_015296823.1 / T2Tme^BB22^) (35), one has Rhs repeats fused to a PoNe DNase (WP_015296737.1 / T2Rhs-Nuc^BB22^) (34, 39), and one has no known domain (WP_015313171.1 / T2Unkwn^BB22^). We could not predict the activity of the latter using sequence (BLAST (40) and HHpred (41)) and structure (AlphaFold2 structure prediction (42, 43) followed by DALI server (44)) analyses. Notably, all five putative effectors are encoded outside of the main T6SS2 gene cluster.

**Fig. 1.**
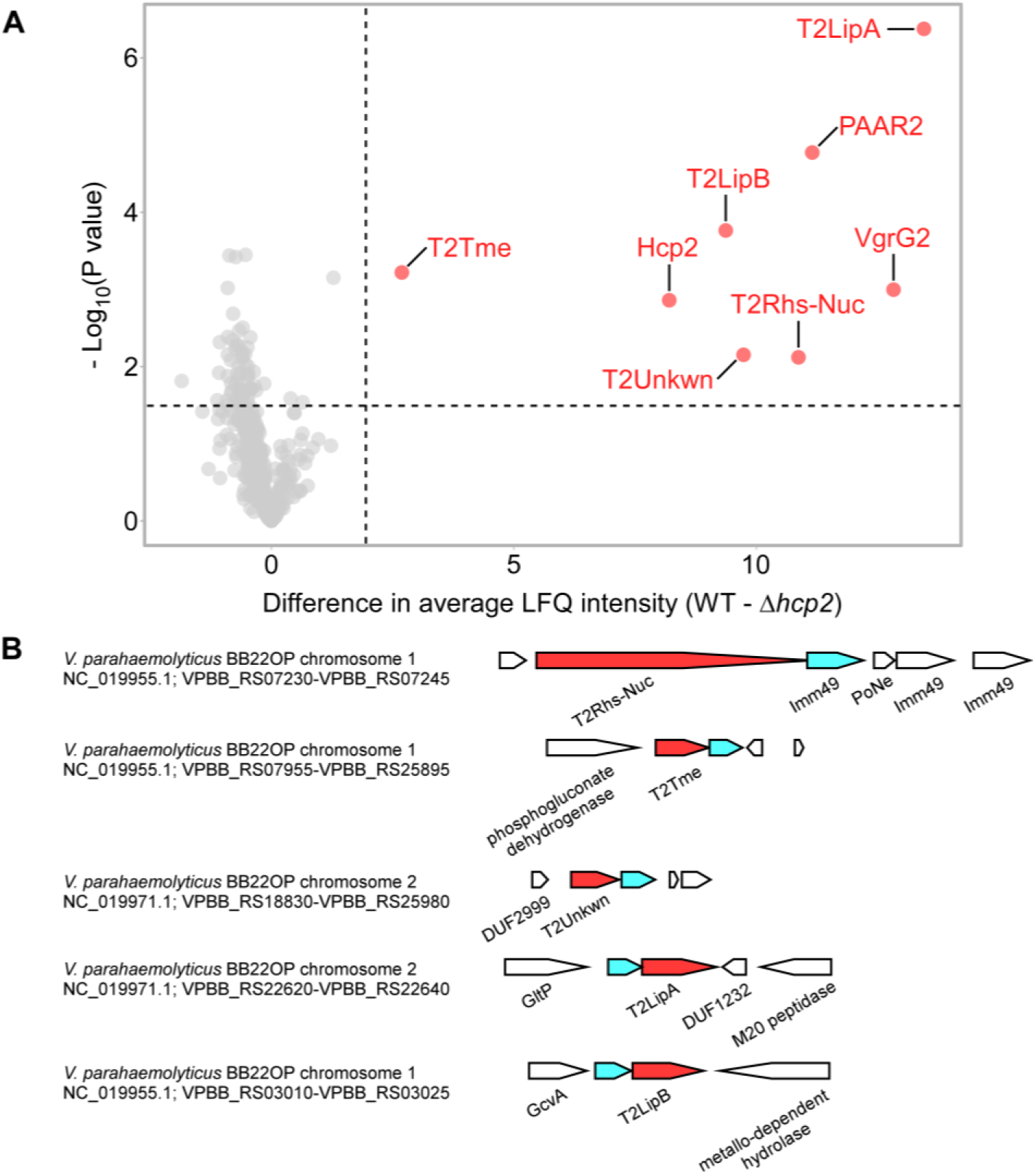
The T6SS2 secretome of *V. parahaemolyticus* strain BB22OP. **(A)** A volcano plot summarizing the comparative proteomics of proteins identified in the medium of wild type (WT) and T6SS2^-^ (Δ*hcp2*) *V. parahaemolyticus* BB22OP strains using label-free quantification. The average difference in signal intensities between the WT strain and the *Δhcp2* strain is plotted against the -Log10 of Student’s *t-*test P values (*n* = 3 biological replicates). Proteins that were significantly more abundant in the secretome of the WT strain (difference in the average LFQ intensities > 2; P value < 0.03; with a minimum of 5 Razor unique peptides) are denoted in red and annotated. **(B)** Schematic representation of genome neighborhoods for non-structural T6SS2-secreted proteins identified in (A). Predicted secreted effectors are denoted in red; predicted neighboring immunity genes are denotes in cyan. Arrows indicate the direction of transcription, and the names of encoded proteins or domains are denoted below. The RefSeq GenBank accession number and the locus tag range are provided.

**Table 1.** Predicted effectors secreted by *V. parahaemolyticus* T6SS2.

| *V. parahaemolyticus* strain | Predicted effector name | Protein accession number | Locus | Known/predicted activity or domain *,^,# |
| --- | --- | --- | --- | --- |
| BB22OP | T2Rhs-Nuc<sup>BB22</sup> | WP_015296737.1 | VPBB_RS07235 | Rhs repeats + PoNe * |
|  | T2Tme<sup>BB22</sup> | WP_015296823.1 | VPBB_RS07950 | Tme ^ |
|  | T2Unkwn<sup>BB22</sup> | WP_015313171.1 | VPBB_RS18835 | Unknown |
|  | T2LipA<sup>BB22</sup> | WP_015313641.1 | VPBB_RS22630 | Lipase_3 * |
|  | T2LipB<sup>BB22</sup> | WP_015296300.1 | VPBB_RS03020 | Lipase_3 * |
| RIMD 2210633 | T2Rhs-Nuc<sup>RIMD</sup> | BAC59780.1 / WP_005479434.1 | VP1517 | Rhs repeats + HNH nuclease * |
| | T2Hydro<sup>RIMD</sup> | BAC61690.1 | VPA0347 | $\alpha/\beta$ hydrolase # |
|  | T2LipB<sup>RIMD</sup> | BAC58889.1 / WP_005483054.1 | VP0626 | Lipase_3 * |

\* Predicted by the NCBI conserved Domain Database; ^ predicted by sequence homology; # predicted by HHpred.

### Identifying the T6SS2 secretome in *V. parahaemolyticus* RIMD 2210633

Next, we wanted to determine whether the T6SS2 secretome differs between *V. parahaemolyticus* strains. We previously reported that T6SS2 in *V. parahaemolyticus* RIMD 2210633 is inactive under warm, marine-like conditions (21). However, Metzger et al. recently found that T6SS2 in this strain plays a role in interbacterial competition when bacteria are grown in low salt concentrations (i.e., in LB media) and upon the over-expression of TfoX, a regulator of T6SS and competence in vibrios (36). This observation suggests that the effectors secreted by this strain are also antibacterial. To confirm this report, we monitored the effect of TfoX over-expression on T6SS2 activity in strain RIMD 2210633. Indeed, we found that upon the arabinose-inducible expression of TfoX, T6SS2 in strain RIMD 2210633 was induced, as manifested by the elevated expression and secretion of Hcp2, a hallmark secreted component of the system (13)(16) (**Fig. 2A**). This activation was independent of surface sensing, which was induced by the addition of phenamil, an inhibitor of the polar flagella motor (21). Moreover, we confirmed that the over-expression of TfoX increased the antibacterial activity of T6SS2, compared to a strain harboring an empty plasmid, as manifested by the reduced viability of *V. natriegens* prey cells during competition on solid agar plates (**Fig. 2B**). Inactivation of T6SS2 by deleting *hcp2* abolished the killing of *V. natriegens* prey, indicating that the TfoX-induced killing was mediated by T6SS2. Notably, since T6SS1 also contributed to bacterial killing under the assay conditions (**Supplementary Fig. S1**), we used a T6SS1^-^ (*Δhcp1*) parental strain for these competition assays.

**Fig. 2.**
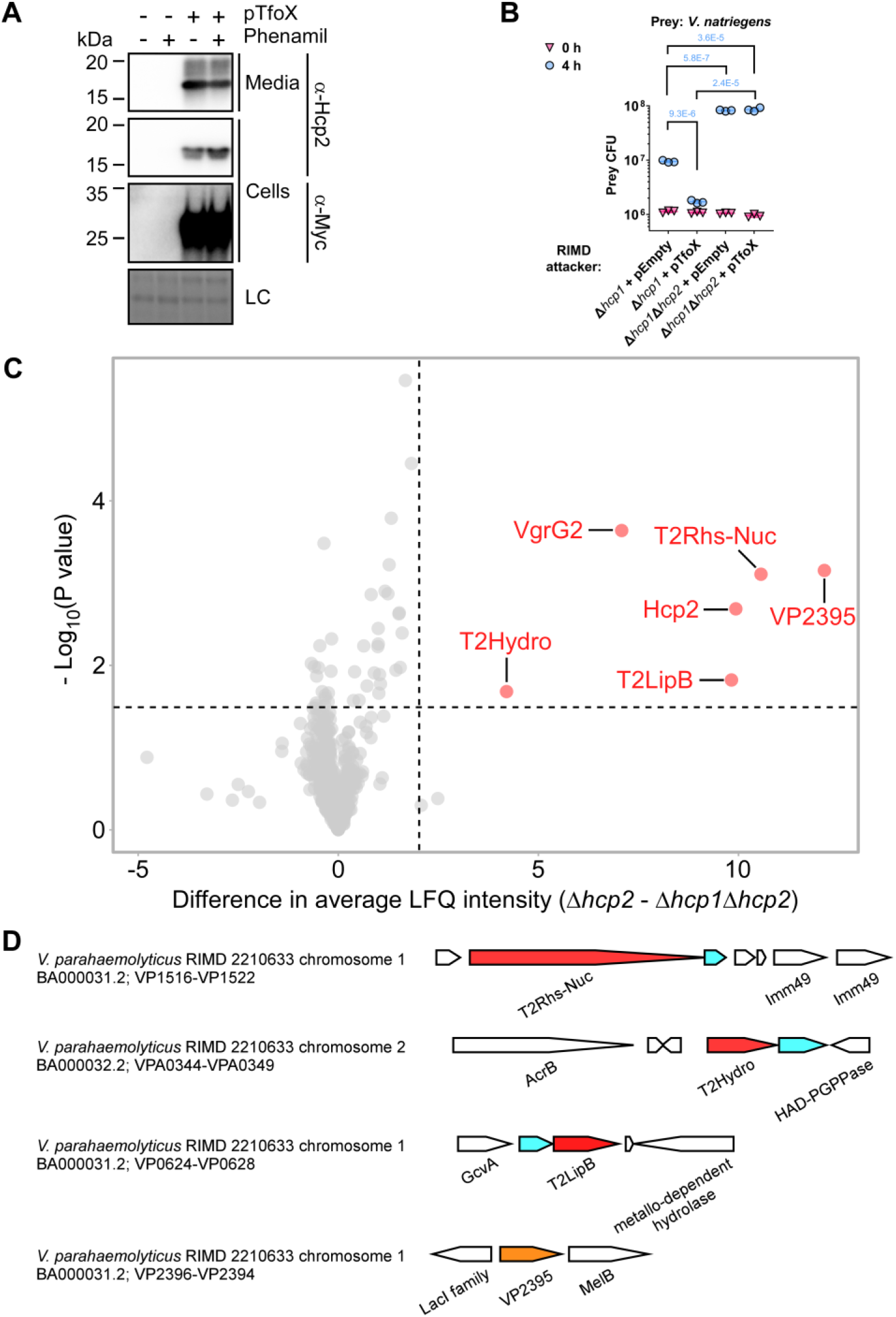
The T6SS2 secretome of *V. parahaemolyticus* strain RIMD 2210633. **(A)** Expression (cells) and secretion (media) of Hcp2 from the *V. parahaemolyticus* RIMD 2210633 strain containing an empty plasmid (pTfoX -) or a plasmid for the arabinose-inducible expression of a C-terminal Myc-His tagged TfoX (pTfoX +). Samples were grown in LB media supplemented with kanamycin to maintain the plasmids, and in the presence (+) or absence (-) of phenamil at 30°C. Loading control (LC) is shown for total protein lysates. **(B)** Viability counts (colony forming units [CFU]) of *V. natriegens* prey strain before (0 h) and after (4 h) co-incubation with the indicated *V. parahaemolyticus* RIMD 2210633 attacker strain carrying an empty plasmid (pEmpty) or a plasmid for the arabinose-inducible expression of TfoX (pTfoX) on LB agar plates supplemented with 0.1% (wt/vol) L-arabinose to induce expression from plasmids. The statistical significance between samples at the 4 h time point was calculated using an unpaired, two-tailed Student’s *t*-test. Data are shown as the mean ± SD; *n* = 3. **(C)** A volcano plot summarizing the comparative proteomics of proteins identified in the medium of the T6SS2^+^ (*Δhcp1*) and T6SS2^-^ (Δ*hcp1*Δ*hcp2*) *V. parahaemolyticus* RIMD 2210633 strains, expressing Tfox from a plasmid, using label-free quantification. The average difference in the signal intensities between the WT strain and the *Δhcp2* strain is plotted against the -Log10 of Student’s *t-*test P values (*n* = 3 biological replicates). Proteins that were significantly more abundant in the secretome of the WT strain (difference in the average LFQ intensities > 2; P value < 0.03; with a minimum of 5 Razor unique peptides) are denoted in red and annotated. **(D)** Schematic representation of genome neighborhoods for non-structural T6SS2-secreted proteins identified in (C). Predicted secreted effectors are denoted in red; predicted neighboring immunity genes are denoted in cyan; *vp2395*, which is not predicted to be an effector, is denoted in orange. Arrows indicate the direction of transcription, and the names of encoded proteins or domains are denoted below. The RefSeq GenBank accession number and the locus tag range are provided.

To reveal the secretome of the RIMD 2210633 T6SS2, we employed the comparative proteomics approach described above for strain BB22OP. To this end, we compared the proteins secreted by a strain with a functional T6SS2 (T6SS2^+^; Δ*hcp1*) with those secreted by a strain in which we inactivated T6SS2 (T6SS2^-^; *Δhcp1Δhcp2);* notably, TfoX was over-expressed in these strains to induce T6SS2. We identified six proteins that were significantly enriched in the secretome of the T6SS2^+^ (**Fig. 2C**, **Table 1,** and **Supplementary Dataset S2**). These proteins include two of the secreted structural components of the T6SS tube-spike complex, Hcp2 and VgrG2 (13), as well as four additional proteins. Three of the additional proteins are predicted to be antibacterial effectors, since they are encoded next to a predicted cognate immunity protein. These include two that are orthologs of proteins identified in the T6SS2 secretome of strain BB22OP and are encoded in the same synteny (i.e., T2Rhs-Nuc^RIMD^ / BAC59780.1 and T2LipB^RIMD^ / BAC58889.1), and another that has a predicted α/β-hydrolase domain (i.e., T2Hydro^RIMD^ / BAC61690.1) (**Fig. 1D**). All three proteins are encoded outside of the T6SS2 gene cluster. The fourth protein, VP2395 (BAC60658.1 / WP_005456695.1), is a predicted cellulase; it is encoded next to genes encoding proteins involved in sugar metabolism. Since *vp2395* does not neighbor a gene that is likely to encode a cognate immunity protein, and given its genomic neighborhood, we did not predict that VP2395 is an antibacterial effector and we therefore omitted it from subsequent analyses. Taken together, these results indicate that the T2Rhs-Nuc orthologs and the T2LipB orthologs are secreted by T6SS2 of both strains, whereas each strain has an additional, different set of proteins secreted by T6SS2.

### Validating T6SS2 effector and immunity pairs

After identifying the T6SS2 secretome in two *V. parahaemolyticus* strains, we set out to determine whether the T6SS2-secreted proteins that we identified in our comparative proteomic analyses are antibacterial T6SS effectors, and whether their downstream- or upstream-encoded proteins serve as cognate immunity proteins. To this end, we generated strains in which we deleted the genes encoding the predicted effectors together with their predicted adjacent immunity genes, and used them as prey strains in self-competition assays. Since the activity of T6SS2 in strain BB22OP was comparable in low (LB, 1% [wt/vol] NaCl) and high (MLB,3% [wt/vol] NaCl) salt media (**Supplementary Fig. S2**), we used LB media for all subsequent competition assays for both BB22OP and the RIMD 2210633 strains.

As shown in **Fig. 3A-D**, deletion of the genes encoding the predicted *V. parahaemolyticus* BB22OP effectors T2Rhs-Nuc^BB22^, T2Tme^BB22^, T2Unkwn^BB22^, and T2LipA^BB22^, together with their neighboring predicted immunity genes (**Fig. 1B**), resulted in prey strains that were sensitive to an attack by their parental wild-type strain. The sensitivity was dependent on a functional T6SS2 in the attacker strain and on the presence of the predicted effector, since deletion of either *hcp2* or the predicted effector alleviated this toxicity. Moreover, expression of the respective, predicted immunity protein from a plasmid in the sensitive prey strain protected it from this T6SS2-mediated attack. Taken together, these results confirm that T2Rhs-Nuc^BB22^, T2Tme^BB22^, T2Unkwn^BB22^, and T2LipA^BB22^ are *bona fide* T6SS2 effectors, and that their neighboring genes encode for their cognate immunity proteins. Surprisingly, although T2LipB^BB22^ and the protein encoded upstream are homologs of the confirmed effector and immunity pair, T2LipA^BB22^-i (36.6% and 27.7% identity, respectively), their deletion did not render the prey strain sensitive to intoxication by a wild-type attacker (**Fig. 3E**). Therefore, we cannot confirm the role of T2LipB^BB22^ as a T6SS2 antibacterial effector at this time.

**Fig. 3.**
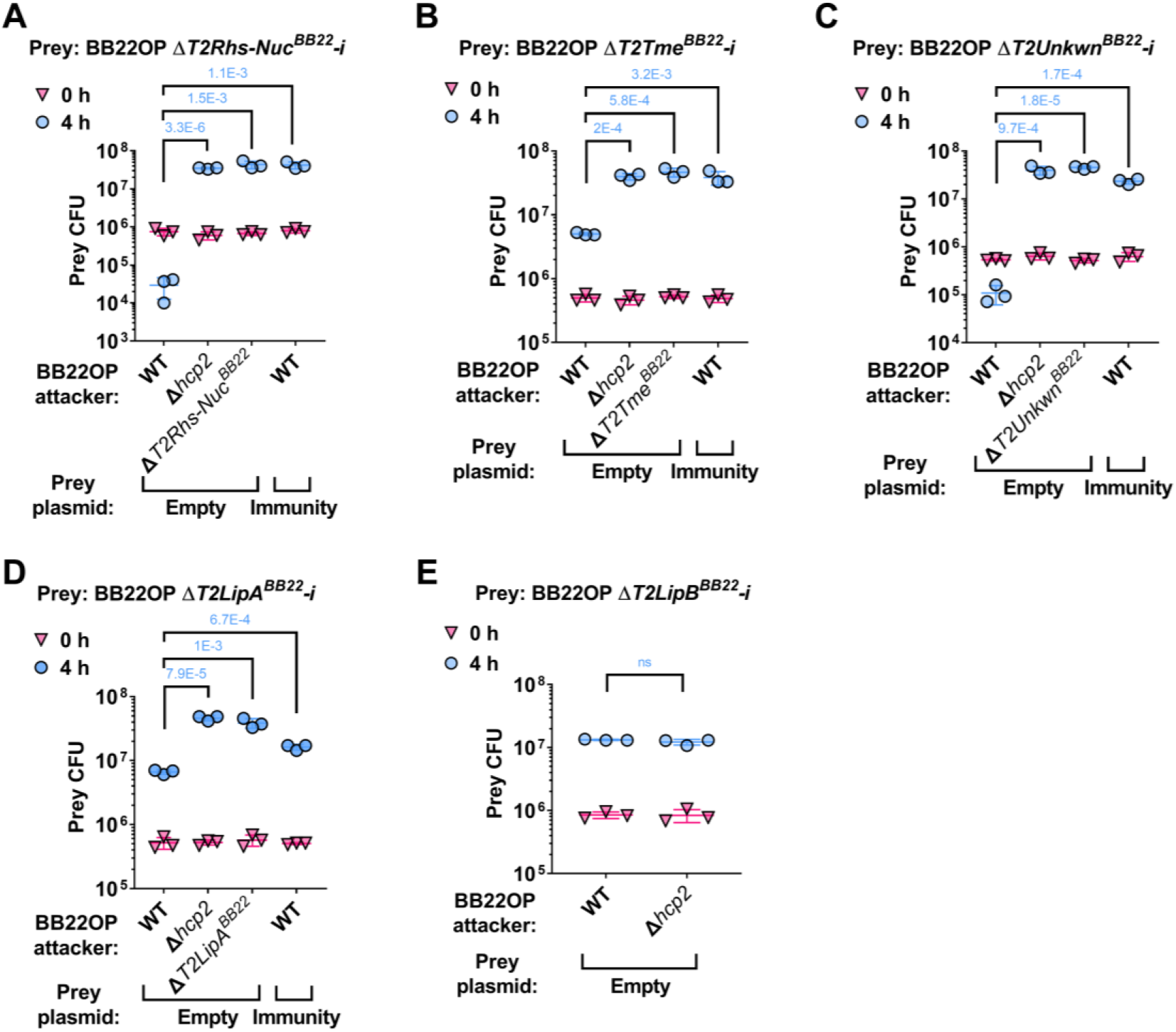
Validation of *V. parahaemolyticus* strain BB22OP T6SS2 effector and immunity pairs. Viability counts (CFU) of the indicated BB22OP derivative prey strains containing a deletion of the predicted effectors T2Rhs-Nuc^BB22^ **(A)**, T2Tme^BB22^ **(B)**, T2Unkwn^BB22^ **(C)**, T2LipA^BB22^ **(D)**, and T2LipB^BB22^ **(E)**, and their neighboring predicted immunity gene (-i) before (0 h) and after (4 h) co-incubation with the indicated *V. parahaemolyticus* BB22OP attacker strains. Prey strains contain either an empty plasmid (Empty) or a plasmid for the arabinose-inducible expression of the predicted immunity protein that was deleted (Immunity). Competitions were performed on LB agar plates supplemented with L-arabinose (0.1% [wt/vol] in A, B, C, and E; 0.01% [wt/vol] in D) to maintain the plasmids and to induce protein expression, respectively. The statistical significance between samples at the 4 h time point was calculated using an unpaired, two-tailed Student’s *t*-test; ns, no significant difference (P > 0.05). Data are shown as the mean ± SD; *n* = 3.

Similarly, deletion of the predicted T6SS2 effectors of *V. parahaemolyticus* RIMD 2210633, T2Rhs-Nuc^RIMD^ and T2Hydro^RIMD^, together with their downstream genes, rendered these prey strains sensitive to an attack by their parental attacker strain (**Fig. 4A-B**). Inactivation of T6SS2 in the attacker strain by deleting *hcp2*, as well as deleting the predicted effector in the attacker strain, alleviated this toxicity. In addition, expression of the downstream gene, predicted to encode the cognate immunity protein, from a plasmid protected the sensitive prey strain from the T6SS2-mediated attack. Taken together, these results confirm that T2Rhs-Nuc^RIMD^ and T2Hydro^RIMD^, together with their downstream genes, constitute *bona fide* T6SS2 effector and immunity pairs. Similar to their orthologs in strain BB22OP (i.e., T2LipB^BB22^ and its predicted immunity; 98% and 99% identity, respectively), deletion of the genes encoding T2LipB^RIMD^ and its predicted immunity protein in strain RIMD 2210633 did not render the prey strain sensitive to a T6SS2-mediated attack by its parental attacker strain (**Fig. 4C**). Therefore, the role of T2LipB^RIMD^ as an antibacterial effector of T6SS2 remains unclear.

**Fig. 4.**
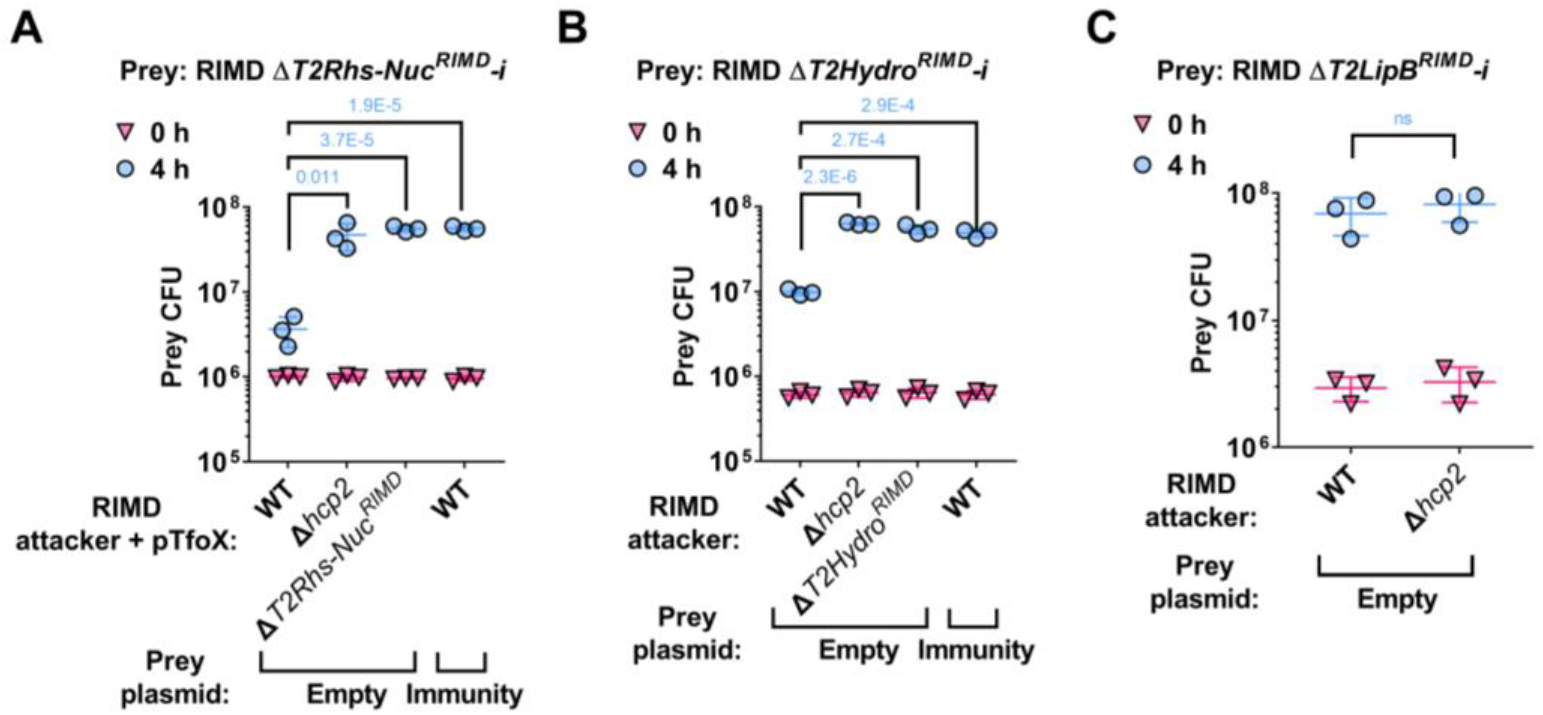
Validation of *V. parahaemolyticus* strain RIMD 2210633 T6SS2 effector and immunity pairs. Viability counts (CFU) of the indicated RIMD 2210633 derivative prey strains containing a deletion of the predicted effectors T2Rhs-Nuc^RIMD^ **(A)**, T2Hydro^RIMD^ **(B)**, and T2LipB^RIMD^ **(C)**, and their neighboring predicted immunity gene (-i) before (0 h) and after (4 h) co-incubation with the indicated *V. parahaemolyticus* RIMD 2210633 attacker strains. Prey strains contain either an empty plasmid (Empty) or a plasmid for the arabinose-inducible expression of the predicted immunity protein that was deleted (Immunity). Competitions were performed on LB agar plates supplemented with 0.1% [wt/vol] L-arabinose to maintain the plasmids and to induce protein expression, respectively. The statistical significance between samples at the 4 h time point was calculated using an unpaired, two-tailed Student’s *t*-test; ns, no significant difference (P > 0.05). Data are shown as the mean ± SD; *n* = 3.

### The start codon of T2Hydro^RIMD^ is misannotated

The newly identified T6SS2 effector in *V. parahaemolyticus* RIMD 2210633, T2Hydro^RIMD^, is annotated with different translational start sites in the two GenBank accessions available on NCBI. In the international nucleotide sequence database collaboration (INSDC) GenBank accession BA000032.2, the effector is annotated as being 467 amino acids long (protein accession number BAC61690.1); in the RefSeq GenBank accession NC_004605.1, it is annotated as a 435 amino acid-long protein (protein accession number WP_021451965.1), with a start site corresponding to methionine 33 in BAC61690.1. However, upon inspection of the protein coverage in our mass spectrometry results, we could not find peptides corresponding to amino acids 1-64 of BAC61690.1. The first amino acid that is covered by peptides identified in the mass spectrometry results corresponds to methionine 65 of BAC61690.1. Based on this observation, we hypothesized that the translation start site of T2Hydro^RIMD^ is misannotated in both the INSDC and RefSeq GenBank accessions, and that the first amino acid of this effector corresponds to methionine 65 in BAC61690.1. This hypothesis was further supported by an analysis performed using the translation start site identification program, Prodigal (45). To determine whether T2Hydro^RIMD^, starting at methionine 65 (T2Hydro^RIMD/M65^), is a functional T6SS2 effector, we set out to test its ability to be delivered by T6SS2 and to mediate interbacterial competition. To this end, we introduced the genes encoding T2Hydro^RIMD/M65^ and its downstream-encoded immunity protein into *V. parahaemolyticus* BB22OP. This strain serves as a surrogate T6SS2-containing attacker, since it has the same T6SS2 as the RIMD 2210633 strain, yet it lacks a T2Hydro^RIMD^ homolog. As shown in **Supplementary Fig. S3**, plasmid-expressed T2Hydro^RIMD/M65^ and its cognate immunity protein enabled a BB22OP surrogate attacker to intoxicate its parental prey (lacking the cognate immunity). This toxicity was dependent on a functional T6SS2 in the surrogate attacker, since it was alleviated upon deletion of *hcp2*. Moreover, expression of the cognate immunity protein from a plasmid in the parental BB22OP prey strain protected it from the T2Hydro^RIMD/M65^-mediated attack. Taken together, these results suggest that the start site of T2Hydro^RIMD^ corresponds to methionine 65 in BAC61690.1.

### T2Rhs-Nuc is conserved and required for T6SS2 activity in *V. parahaemolyticus*

Next, we investigated whether the deletion of the effectors described above affected T6SS2 activity. To this end, we monitored the secretion of Hcp2, a hallmark T6SS-secreted protein, in strains deleted for each of the predicted effectors. Surprisingly, deletion of T2Rhs-Nuc^BB22^ and T2Rhs-Nuc^RIMD^ abolished Hcp2 secretion in their respective strains (**Fig. 5A-B**). These results indicate that T2Rhs-Nuc is not only a T6SS2 effector—it is also required for T6SS2 activity.

Since the investigated T2Rhs-Nuc effectors are not encoded within the T6SS2 gene cluster (**Fig. 1B** and **Fig. 2D**), the observation that they are essential for T6SS2 activity suggests that T2Rhs-Nuc should be conserved in *V. parahaemolyticus* strains. Indeed, we identified T2Rhs-Nuc orthologs in almost all of the complete *V. parahaemolyticus* genomes available on the NCBI RefSeq database (found in 73 out of 75 genomes examined) (**Fig. 5C** and **Supplementary Dataset S3**). This result is in agreement with the requirement of T2Rhs-Nuc for T6SS2 activity.

**Fig. 5.**
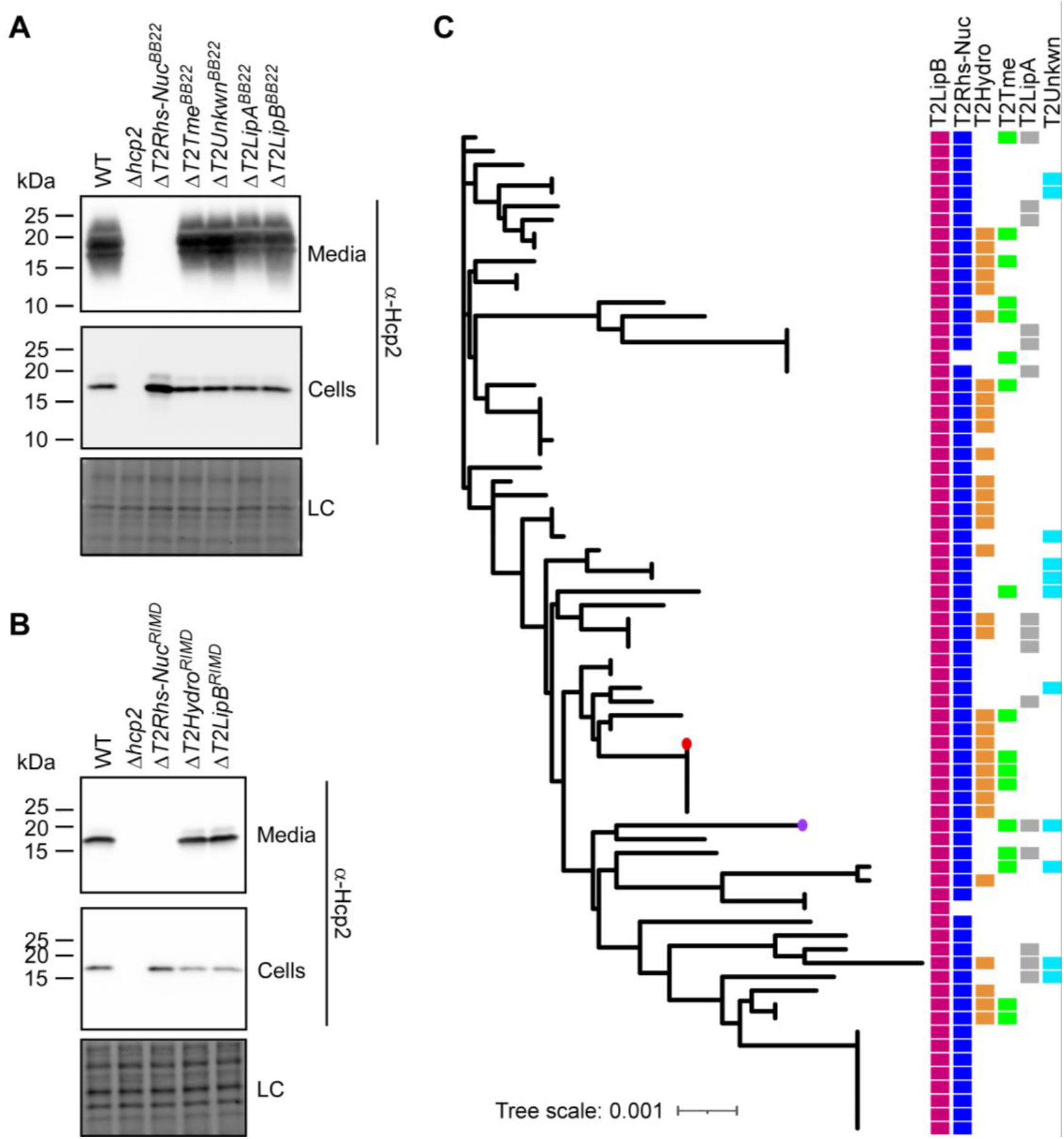
T2Rhs-Nuc is ubiquitous in *V. parahaemolyticus* genomes and is required for T6SS2 activity. **(A-B)** Expression (cells) and secretion (media) of Hcp2 from *V. parahaemolyticus* BB22OP (A) and RIMD 2210633 (B) wild-type (WT) strains or their indicated derivatives containing a deletion in a gene encoding a secreted T6SS2 effector. Samples were grown in LB media at 30°C. Loading control (LC) is shown for total protein lysates. **(C)** Distribution of T6SS2-secreted effectors in complete *V. parahaemolyticus* genomes. The phylogenetic tree was based on DNA sequences of *rpoB* coding for DNA-directed RNA polymerase subunit beta. The evolutionary history was inferred using the neighbor-joining method. *V. parahaemolyticus* strains BB22OP and RIMD 2210633 are denoted by a circle (purple and red, respectively).

Further analysis of complete *V. parahaemolyticus* genomes revealed that, similar to T2Rhs-Nuc, T2LipB homologs are ubiquitous in this species. In contrast, effectors identified in the T6SS2 secretome of only one of the strains that we investigated were encoded only by a subset of the genomes; each genome harbored a different combination of T6SS2-secreted effectors (**Fig. 5C** and **Supplementary Dataset S1**). Taken together, these results indicate that T2RhsNuc and T2LipB constitute the “core” substrates secreted by *V. parahaemolyticus* T6SS2, whereas T2LipA, T2Hydro, T2Tme, and T2Unkwn appear to belong to an accessory T6SS2 effector repertoire.

## Discussion

In a recent report, we found that all strains of the emerging pathogen *V. parahaemolyticus* have a conserved T6SS, named T6SS2 (32), indicating that this system plays a significant role in the life cycle of this bacterium. Although this system has recently been shown to play a role in interbacterial competition (35, 36), its effector repertoire has remained unknown. In this work, we revealed the core and accessory effector repertoires of *V. parahaemolyticus* T6SS2. Notably, all of the identified effectors are encoded outside of the T6SS2 gene cluster.

Comparative proteomic analyses on two *V. parahaemolyticus* strains, BB22OP and RIMD 2210633, revealed two proteins that were secreted by both strains: T2Rhs-Nuc and T2LipB. We found that these two proteins are encoded by nearly all *V. parahaemolyticus* strains for which a complete genome sequence is available on NCBI. Based on these findings, we propose that T2Rhs-Nuc and T2LipB constitute the conserved core of the T6SS2-secreted protein repertoire in *V. parahaemolyticus*.

Furthermore, we found that the conserved T2Rhs-Nuc is required for T6SS2 activity, suggesting that the loading of this effector onto the T6SS serves as a quality control checkpoint to enable T6SS2 delivery. The role of certain effectors as a structural necessity in T6SS assembly was also previously suggested by others (46–48). Donato et al. (48) demonstrated that two Rhs repeat-containing effectors in *Enterobacter cloacae* are required for T6SS-mediated secretion. Notably, in contrast to T2Rhs-Nuc, which is required for T6SS2 activity in *V. parahaemolyticus*, the two *Enterobacter* Rhs repeat-containing proteins are specialized effectors that also contain an N-terminal terminal PAAR domain, which is known to play a structural role in the secreted spike complex by capping the VgrG trimer (49).

Although T2LipB proteins are conserved and secreted by both *V. parahaemolyticus* strains investigated in this work, and even though T2LipB proteins and their predicted upstream-encoded proteins are homologs of the confirmed T6SS2 effector and immunity pair, T2LipA^BB22^-i, we were unable to determine whether T2LipB proteins function as antibacterial effectors. There are two possible explanations of why the deletion of T2LipB2 and its upstream putative immunity gene did not sensitize *V. parahaemolyticus* to attack by a parental strain: (1) T2LipB plays a different role as a secreted T6SS2 protein and it is not an antibacterial effector; (2) *V. parahaemolyticus* is not sensitive to intoxication by T2LipB due to the presence of a yet-to-be-identified immunity protein or because of an immunity protein-independent defense mechanism. Future work focusing on T2LipB is required to determine its role as a T6SS2-secreted protein.

In addition to the two conserved secreted proteins, T2Rhs-Nuc and T2LipB, we identified four effectors that were secreted by T6SS2 of either strain BB22OP or strain RIMD 2210633: T2LipA, T2Tme, T2Unkwn, and T2Hydro. Since we found that these effectors are differentially distributed among *V. parahaemolyticus* strains, we concluded that they represent at least a subset of the T6SS2 accessory effector repertoire. We hypothesize that other *V. parahaemolyticus* strains carry additional effectors that belong to the T6SS2 accessory effector repertoire.

In conclusion, we identified several effectors secreted by the conserved *V. parahaemolyticus* T6SS2, and we found that one of the conserved effectors, T2Rhs-Nuc, plays another role as a quality control checkpoint that is required for T6SS2 activity. These results confirm the predicted role of this T6SS in interbacterial competitions, and enlarge the repertoire of known T6SS effectors. We find T2Unkwn of special interest, since it does not resemble any previously described toxin; future work may reveal its mechanism of action and target.

## Materials and Methods

### Strains and Media

For a complete list of strains used in this study, see **Supplementary Table S1**. *Escherichia coli* strain DH5α (*λ-pir*) was grown in lysogeny broth (LB; containing 1% [wt/vol] NaCl) or on LB agar (1.5% [wt/vol]) plates at 37°C, or at 30°C when harboring effector expression plasmids. Media were supplemented with chloramphenicol (10 μg/ml), kanamycin (30 μg/ml), and gentamycin (50 μg/ml) when needed to maintain plasmids. Glucose (0.4% [wt/vol]) was added to repress protein expression from the arabinose-inducible promoter, *Pbad*. To induce expression from P*bad*, L-arabinose was added to the media at 0.01 or 0.1% (wt/vol), as indicated. *Vibrio parahaemolyticus* strains BB22OP, RIMD 2210633, and their derivatives, as well as *Vibrio natriegens* ATCC 14048, were grown in Marine Lysogeny Broth (MLB; LB containing 3% [wt/vol] NaCl) and on Marine Minimal Media (MMM) agar plates (1.5% [wt/vol] agar, 2% [wt/vol] NaCl, 0.4% [wt/vol] galactose, 5 mM MgSO_4_, 7 mM K_2_SO_4_, 77 mM K_2_HPO_4_, 35 mM KH_2_PO_4_, and 2 mM NH4Cl) at 30°C. Media were supplemented with chloramphenicol (10 μg/ml), kanamycin (250 μg/ml), or gentamycin (50 μg/ml) to maintain plasmids. To induce expression from P*bad*, L-arabinose was added to the media at 0.01 or 0.1% (wt/vol), as indicated.

### Plasmid construction

For a complete list of plasmids used in this study, see **Supplementary Table S2**. For expression in bacteria, the coding sequences (CDS) of the genes of interest were PCR amplified from the respective genomic DNA of the encoding bacterium. Next, amplicons were inserted into the multiple cloning site (MCS) of pBAD^K^/Myc-His, pBAD33.1^F^ or their derivatives using the Gibson assembly method (50). Plasmids were introduced into *E. coli* DH5α (*λ-pir*) by electroporation, and into vibrios via conjugation. Transconjugants were selected on MMM agar plates supplemented with the appropriate antibiotics to select clones containing the desired plasmids.

### Construction of deletion strains

The construction of in-frame deletions in *V. parahaemolyticus* strains was described previously (21, 34). Briefly, 1 kb sequences upstream and downstream of each gene or operon to be deleted were cloned into pDM4, a Cm^R^ OriR6K suicide plasmid. The pDM4 constructs were transformed into *E. coli* DH5α (*λ-piŕ*) by electroporation, and then transferred into vibrios via conjugation. Transconjugants were selected on MMM agar plates supplemented with chloramphenicol, and then counter-selected on MMM agar plates containing 15% (wt/vol) sucrose for loss of the sacB-containing plasmid. Deletions were further confirmed by PCR.

### Bacterial competition assays

Bacterial competition assays were performed as previously described (21), with minor modifications. Briefly, cultures of the indicated attacker and prey strains were grown overnight. Bacterial cultures were then normalized to OD_600_ = 0.5 and mixed at a 10:1 (attacker:prey) ratio in triplicate. Next, the mixtures were spotted (25 μl) on LB or MLB agar plates supplemented with 0.01 or 0.1% (wt/vol) L-arabinose, as indicated, and incubated for 4 h at 30°C. The colony-forming units (CFU) of the prey strains were determined at the 0 and 4 h time points by counting 10-fold serial dilutions plated on MMM agar plates, supplemented with an appropriate antibiotic to select for prey colony growth. The experiments were performed at least three times with similar results. Results from a representative experiment are shown.

### Hcp2 secretion assays

Hcp2 secretion assays were performed as previously described (21), with minor modifications. Briefly, *Vibrio* strains were grown overnight in MLB broth supplemented with antibiotics to maintain plasmids when needed. Bacterial cultures were then normalized to an OD_600_ of 0.18 (BB22OP) or 0.9 (RIMD 2210633) in 5 ml LB broth supplemented with appropriate antibiotics and 0.1% (wt/vol) L-arabinose when expression from an arabinose-inducible plasmid was required. Bacterial cultures were incubated with constant shaking (220 rpm) at 30°C for 5 h. For expression fractions (cells), cells equivalent to 1 OD_600_ units were collected, and cell pellets were resuspended in 100 μl of 2x Tris-glycine SDS sample buffer (Novex, Life Sciences). For secretion fractions (media), supernatant volumes equivalent to 10 OD_600_ units were filtered (0.22 μm), and proteins were precipitated using the deoxycholate and trichloroacetic acid method (51). The precipitated proteins were washed twice with cold acetone, and then air-dried before resuspension in 20 μl of 100 mM Tris-Cl (pH = 8.0) and 20 μl of 2X protein sample buffer. Next, samples were incubated at 95°C for 5 or 10 min and then resolved on TGX Stain-free gel (Bio-Rad). The proteins were transferred onto 0.2 μm nitrocellulose membranes using Trans-Blot Turbo Transfer (Bio-Rad) according to the manufacturer’s protocol. Membranes were then immunoblotted with custom-made α-Hcp2 (52) or α-Myc (Santa Cruz Biotechnologies, sc-40) antibodies at 1:1000 dilution. Protein signals were visualized in a Fusion FX6 imaging system (Vilber Lourmat) using enhanced chemiluminescence (ECL) reagents. The experiments were performed at least three times with similar results. Results from a representative experiment are shown.

### Mass spectrometry analyses

Sample preparation for mass spectrometry was performed as described in the “Hcp2 secretion assays” section, with minor modifications (*V. parahaemolyticus* BB22OP strains were grown in MLB media, and *V. parahaemolyticus* RIMD2210633 strains were grown in LB media). After the acetone wash step, samples were shipped to the Smoler Proteomics Center at the Technion for analysis.

Precipitated proteins were washed 3 times in cold 80% (v/v) acetone and incubated for 15 min at −20°C, followed by centrifugation at 16,000 x *g* for 10 min at 4°C. The protein pellets were then resuspended and incubated at 60°C for 30 min in reducing urea buffer (8 M urea, 100 mM ammonium bicarbonate, and 3 mM DTT for BB22OP samples; 9 M urea, 400 mM ammonium bicarbonate, and 10 mM DTT for RIMD 221063 samples). The proteins were then modified with iodoacetamide (45 and 35 mM for the RIMD 2210633 and BB22OP samples, respectively) in 100 mM ammonium bicarbonate for 30 min at room temperature in the dark. Then, 10 μg of protein were digested overnight at 37°C in 2 or 1.5 M urea (for BB22OP and RIMD 221063 samples, respectively) and 25 mM ammonium bicarbonate with modified trypsin (Promega), in a 1:50 (M/M) enzyme-to-substrate ratio. For the RIMD 221063 samples, an additional trypsinization step was performed for 4 h. The tryptic peptides were acidified by adding 1% formic acid and desalted using C18 tips (homemade stage tips), then dried and re-suspended in 0.1% Formic acid. The resulting tryptic peptides were resolved by reverse-phase chromatography on 0.075 × 250-mm or 0.075 × 300-mm (for BB22OP and RIMD 2210633 samples, respectively) fused silica capillaries (J&W) packed with Reprosil reversed phase material (Dr Maisch GmbH, Germany). The peptides were eluted with a linear 60 min gradient of 5 to 28%, 15 min gradient of 28 to 95%, and 15 min at 95% acetonitrile with 0.1% formic acid in water at a flow rate of 0.15 μl/min. Mass spectrometry was performed using a Q Exactive plus mass spectrometer (Thermo) in a positive mode using a repetitively full MS scan followed by high collision dissociation (HCD) of the 10 most dominant ions selected from the first MS scan.

The mass spectrometry data were analyzed using MaxQuant software 1.5.2.8 for peak picking and identification using the Andromeda search engine (53) against the relevant *V. parahaemolyticus* strain from the Uniprot database with mass tolerance of 6 ppm for the precursor masses and 20 ppm for the fragment ions. For the BB22OP samples, oxidation on methionine was accepted as variable modifications, and carbamidomethyl on cysteine was accepted as static modifications. The minimal peptide length was set to six amino acids; a maximum of two miscleavages was allowed. For the RIMD 2210633 samples, oxidation on methionine and protein N-terminus acetylation were accepted as variable modifications, and carbamidomethyl on cysteine was accepted as static modifications. The minimal peptide length was set to 7 amino acids; a maximum of two miscleavages was allowed. For the BB22OP samples, the minimum number of samples identified per protein was set to 2. Peptide-level and protein-level false discovery rates (FDRs) were filtered to 1% using the target-decoy strategy. Protein tables were filtered to eliminate the identifications from the reverse database and common contaminants and single peptide identifications. The data were quantified by label-free analysis using the same software, based on extracted ion currents (XICs) of peptides enabling quantitation from each LC/MS run for each peptide identified in any of the samples. Statistical analysis of the identification and quantization results was done using Perseus 1.6.7.0 software (54). Intensity data were transformed to log2. Missing values were replaced with 18 (on the logarithmic scale), which corresponds to the lowest intensity that was detected. A Student’s *t*-test with Permutation-based FDR (with 250 randomization, threshold value = 0.05) was performed.

The mass spectrometry proteomics data have been deposited in the ProteomeXchange Consortium via the PRIDE (55) partner repository with the dataset identifiers PXD037864 for strain BB22OP, and PXD037980 for strain RIMD 2210633.

### Identifying effector homologs in *V. parahaemolyticus* genomes

A local database containing the RefSeq bacterial nucleotide and protein sequences was generated (last updated on June 11, 2022). BLASTP was employed to identify homologs of the T6SS2 effectors in *V. parahaemolyticus* complete genomes. The amino acid sequences of T2Rhs-Nuc^RIMD^ (WP_005479434.1), T2Hydro^RIMD^ (BAC61690.1, amino acids 65-467), T2LipA^BB22^ (WP_015313641.1), T2Tme^BB22^ (WP_015296823.1), T2Unkwn^BB22^ (WP_015313171.1), and T2LipB^BB22^ (WP_015296300.1) were used as queries. The E-value threshold was set to 10^-12^ and the coverage was set to 70%, based on the length of the query sequences.

### Constructing a phylogenetic tree

The nucleotide sequences of *rpoB* were retrieved from the local RefSeq database (partial and pseudogene sequences were removed). Phylogenetic analyses of bacterial genomes were conducted using the MAFFT 7 server (mafft.cbrc.jp/alignment/server/). Multiple sequence alignment was generated using MAFFT v7 FFT-NS-i (56, 57). The evolutionary history of *V. parahaemolyticus* genomes was inferred using the neighbor-joining method (58) with the Jukes-Cantor substitution model (JC69). The analysis included 73 nucleotide sequences and 4,029 conserved sites.

## Data Availability Statement

The authors confirm that the data supporting the findings of this study are available within the article and its supplementary material. The mass spectrometry raw data files were deposited in ProteomeXchange under the accession numbers indicated in the Materials and Methods section.

## Acknowledgments

This project received funding from the Israel Science Foundation (grant no. 920/17 to D. Salomon, and grant no. 1362/21 to D. Salomon and E. Bosis). We thank members of the Salomon and Bosis labs for helpful discussions and suggestions. We also thank the Smoler Proteomics Center at the Technion for performing and analyzing the mass spectrometry data. This work was performed in partial fulfillment of the requirements for a PhD degree for D. Tchelet at the Sackler Faculty of Medicine, Tel Aviv University.

## Author Contributions

D. Tchelet: conceptualization, investigation, methodology, and writing—original draft.

K. Keppel: investigation and methodology.

E. Bosis: conceptualization, investigation, methodology, funding acquisition, and writing— original draft.

D. Salomon: conceptualization, supervision, funding acquisition, investigation, methodology, and writing—original draft.

## Conflict of Interest

The authors declare that they have no conflict of interest.

## Notes

### Competing Interest Statement

The authors have declared no competing interest.

