## Supplementary Information for "*Vibrio parahaemolyticus* T6SS2 effector repertoires"

**Supplementary Figures S1-S3**

**Supplementary Datasets S1-S3**

**Supplementary Tables S1-S2**

**Supplementary References**

### Supplementary Figures

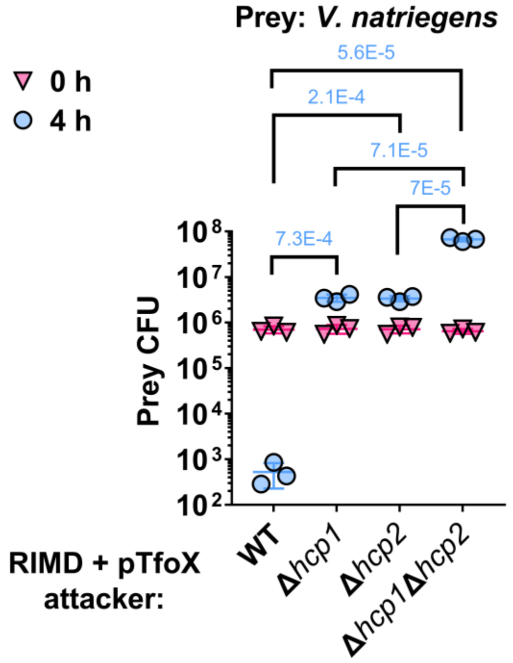


**Supplementary Fig. S1. T6SS1 and T6SS2 play a role in interbacterial competition in *V. parahaemolyticus* strain RIMD 2210633.** Viability counts (CFU) of the *V. natriegens* prey strain before (0 h) and after (4 h) co-incubation with the indicated *V. parahaemolyticus* RIMD 2210633 attacker strains carrying a plasmid for the arabinose-inducible expression of TfoX (pTfoX) on LB agar plates supplemented with 0.1% (wt/vol) L-arabinose to induce expression from plasmids. The statistical significance between samples at the 4 h time point was calculated using an unpaired, two-tailed Student's *t*-test. Data are shown as the mean ± SD; *n* = 3.

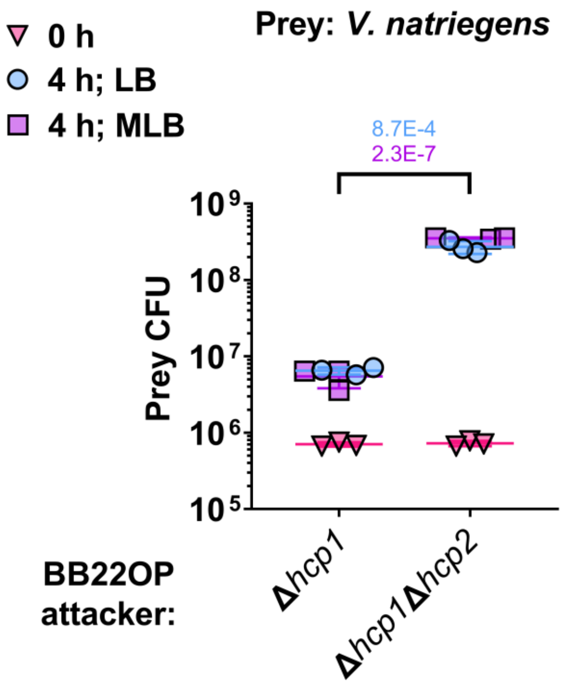


**Supplementary Fig. S2. The activity of *V. parahaemolyticus* BB22OP T6SS2 is comparable in LB and MLB media.** Viability counts (CFU) of *V. natriegens* prey strain before (0 h) and after (4 h) co-incubation with the indicated *V. parahaemolyticus* BB22OP attacker strains on LB or MLB agar plates. The statistical significance between samples at the 4 h time point was calculated using an unpaired, two-tailed Student's *t*-test. Data are shown as the mean ± SD; $n = 3$.

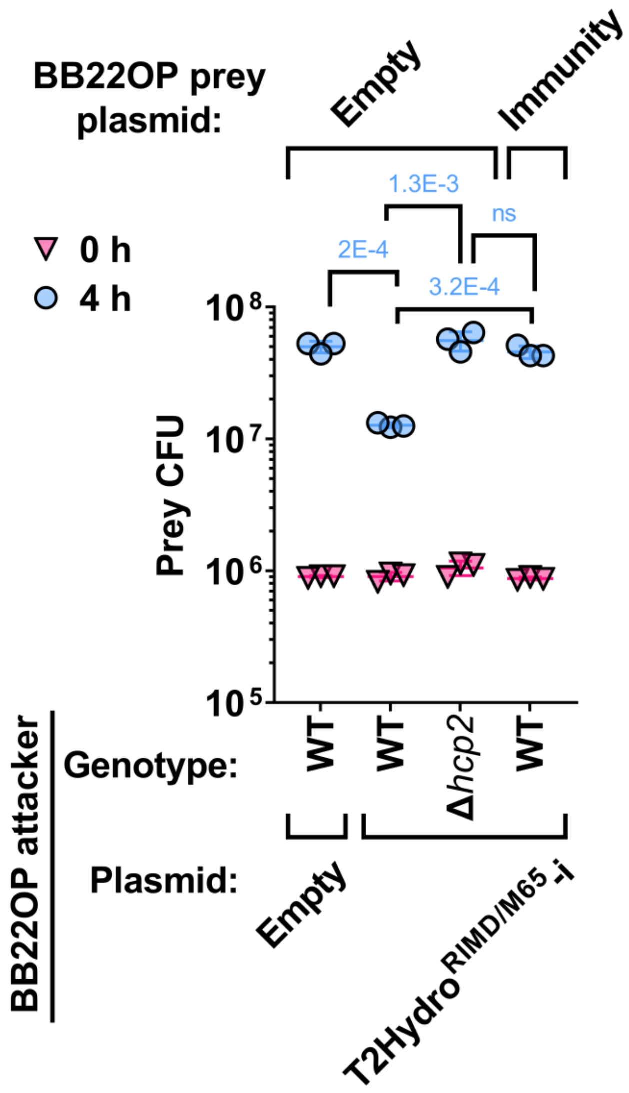


**Supplementary Fig. S3. T2Hydro$^{RIMD/M65}$ is a functional T6SS2 effector.** Viability counts (CFU) of *V. parahaemolyticus* BB22OP prey strains containing either an empty plasmid (Empty) or a plasmid for the arabinose-inducible expression of the T2Hydro$^{RIMD}$ cognate protein (Immunity) before (0 h) and after (4 h) co-incubation with the indicated *V. parahaemolyticus* BB22OP attacker strains carrying an empty plasmid or a plasmid for the arabinose-inducible expression of T2Hydro$^{RIMD/M65}$ and its downstream-encoded immunity protein (T2Hydro$^{RIMD/M65}$-i) on LB agar plates supplemented with 0.1% (wt/vol) L-arabinose to induce expression from plasmids. The statistical significance between samples at the 4 h time point was calculated using an unpaired, two-tailed Student's *t*-test; ns, no significant difference ($P > 0.05$). Data are shown as the mean ± SD; $n = 3$.

### Supplementary Datasets

**Supplementary Dataset S1. Mass spectrometry results for the *V. parahaemolyticus* BB22OP samples.**

**Supplementary Dataset S2. Mass spectrometry results for the *V. parahaemolyticus* RIMD 2210633 samples.**

**Supplementary Dataset S3. The distribution of T6SS2 effectors in complete *V. parahaemolyticus* genomes.**

### Supplementary Tables

#### Supplementary Table S1. A list of bacterial strains used in this work.

| Strain name | Genotype | Comments | Source |
|---|---|---|---|
| *Vibrio parahaemolyticus* RIMD 2210633 | Wild type | Used for generating deletion strains, Hcp2 secretion assays, and as an attacker in competition assays | Obtained from Kim Orth |
| *Vibrio parahaemolyticus* BB22OP | Wild type | Used for generating deletion strains, Hcp2 secretion assays, proteomic analysis and as an attacker in competition assays | Obtained from Kim Orth |
| *Vibrio natriegens* ATCC 14048 | Wild type | Used as prey in competition assays | ATCC collection |
| *Vibrio parahaemolyticus* RIMD 2210633 Δ*hcp1* | Δ*vp1393* | RIMD 2210633 derivative containing an in-frame deletion of *vp1393;* used in proteomic analysis and as an attacker in competition assays | (1) |
| *Vibrio parahaemolyticus* RIMD 2210633 Δ*hcp2* | Δ*vpa1027* | RIMD 2210633 derivative containing an in-frame deletion of *vpa1027*; used in secretion assays and as an attacker in competition assays | This study |
| *Vibrio parahaemolyticus* RIMD 2210633 Δ*hcp1*Δ*hcp2* | Δ*vp1393*Δ*vpa1027* | RIMD 2210633 derivative containing an in-frame deletion of *vp1393* and *vpa1027*; used in proteomic analysis and as an attacker in competition assays | This study |
| *Vibrio parahaemolyticus* RIMD 2210633 Δ*T2Rhs-Nuc*$^{RIMD}$ | Δ*vp1517* | RIMD 2210633 derivative containing an in-frame deletion of *vp1517*; used in Hcp2 secretion assays and as an attacker in competition assays | This study |

| | | | |
|---|---|---|---|
| *Vibrio parahaemolyticus* RIMD 2210633 Δ*T2Rhs-Nuc$^{RIMD}$-i* | Δ*vp1517/8* | RIMD 2210633 derivative containing an in-frame deletion of *vp1517* and *vp1518*; used as prey in competition assays | This study |
| *Vibrio parahaemolyticus* RIMD 2210633 Δ*T2Hydro$^{RIMD}$* | Δ*vpa0347* | RIMD 2210633 derivative containing an in-frame deletion of *vpa0347*; used in Hcp2 secretion assays and as an attacker in competition assays | This study |
| *Vibrio parahaemolyticus* RIMD 2210633 Δ*T2Hydro$^{RIMD}$-i* | Δ*vpa0347/8* | RIMD 2210633 derivative containing an in-frame deletion of *vpa0347* and *vpa0348*; used as prey in competition assays | This study |
| *Vibrio parahaemolyticus* RIMD 2210633 Δ*T2LipB$^{RIMD}$* | Δ*vp0626* | RIMD 2210633 derivative containing an in-frame deletion of *vp0626*; used in Hcp2 secretion assays | This study |
| *Vibrio parahaemolyticus* RIMD 2210633 Δ*T2LipB$^{RIMD}$-i* | Δ*vp0626/5* | RIMD 2210633 derivative containing an in-frame deletion of *vp0626* and *vp0625*; used as prey in competition assays. | This study |
| *Vibrio parahaemolyticus* BB22OP Δ*hcp1* | Δ*vpbb_rs06665* | BB22OP derivative containing an in-frame deletion of *vpbb_rs06665;* used as an attacker in competition assays | (2) |
| *Vibrio parahaemolyticus* BB22OP Δ*hcp2* | Δ*vpbb_rs19920* | BB22OP derivative containing an in-frame deletion of *vpbb_rs19920;* used in Hcp2 secretion assays, proteomic analysis, and as an attacker in competition assays | (2) |
| *Vibrio parahaemolyticus* BB22OP Δ*hcp1*Δ*hcp2* | Δ*vpbb_rs06665*Δ*vpbb_rs19920* | BB22OP derivative containing an in-frame deletion of Δ*vpbb_rs06665* and Δ*vpbb_rs19920*; used in Hcp2 secretion assays | (2) |

| | | | |
|---|---|---|---|
| | | and as an attacker in competition assays | |
| *Vibrio parahaemolyticus* BB22OP Δ*T2Rhs-Nuc*$^{BB22}$ | Δ*vpbb_rs07235* | BB22OP derivative containing an in-frame deletion of *vpbb_rs07235*; used in Hcp2 secretion assays and as an attacker in competition assays | This study |
| *Vibrio parahaemolyticus* BB22OP Δ*T2Rhs-Nuc*$^{BB22}$*-i* | Δ*vpbb_rs07235/40* | BB22OP derivative containing an in-frame deletion of *vpbb_rs07235* and *vpbb_rs07240*; used as prey in competition assays | This study |
| *Vibrio parahaemolyticus* BB22OP Δ*T2Tme*$^{BB22}$ | Δ*vpbb_rs07950* | BB22OP derivative containing an in-frame deletion of *vpbb_rs07950*; used in Hcp2 secretion assays and as an attacker in competition assays | This study |
| *Vibrio parahaemolyticus* BB22OP Δ*T2Tme*$^{BB22}$*-i* | Δ*vpbb_rs07950/25090* | BB22OP derivative containing an in-frame deletion of *vpbb_rs07950* and *vpbb_rs25090*; used as prey in competition assays | This study |
| *Vibrio parahaemolyticus* BB22OP Δ*T2Unkwn*$^{BB22}$ | Δ*vpbb_rs18835* | BB22OP derivative containing an in-frame deletion of *vpbb_rs18835*; used in Hcp2 secretion assays and as an attacker in competition assays | This study |
| *Vibrio parahaemolyticus* BB22OP Δ*T2Unkwn*$^{BB22}$*-i* | Δ*vpbb_rs18835/40* | BB22OP derivative containing an in-frame deletion of *vpbb_rs18835* and *vpbb_rs18840*; used as prey in competition assays | This study |
| *Vibrio parahaemolyticus* BB22OP Δ*T2LipA*$^{BB22}$ | Δ*vpbb_rs22630* | BB22OP derivative containing an in-frame deletion of *vpbb_rs22630*; used in Hcp2 secretion assays | This study |

| | | | |
|---|---|---|---|
| | | and as an attacker in competition assays | |
| *Vibrio parahaemolyticus* BB22OP Δ*T2LipA$^{BB22}$-i* | Δ*vpbb_rs22630/25* | BB22OP derivative containing an in-frame deletion of *vpbb_rs22630* and *vpbb_rs22625*; used as prey in competition assays | This study |
| *Vibrio parahaemolyticus* BB22OP Δ*T2LipB$^{BB22}$* | Δ*vpbb_rs03020* | BB22OP derivative containing an in-frame deletion of *vpbb_rs22630*; used in Hcp2 secretion assays | This study |
| *Vibrio parahaemolyticus* BB22OP Δ*T2LipB$^{BB22}$-i* | Δ*vpbb_rs03020/15* | BB22OP derivative containing an in-frame deletion of *vpbb_rs03020* and *vpbb_rs03015*; used as prey in competition assays | This study |
| *Escherichia coli* DH5α (*λ-pir*) | K-12 derivative laboratory strain containing λ-pir | Used for plasmid maintenance and cloning | Obtained from Eric V. Stabb |

**Supplementary Table S2. A list of plasmids used in this work.**

| **Plasmid name** | **Description** | **Purpose** | **Source** |
|---|---|---|---|
| pBAD$^{K}$/Myc-His | pBR322 ori-containing plasmid harboring a Kan$^{R}$ cassette, *araC*, and an MCS following a P*bad* promoter | Used for arabinose-inducible expression | (3) |
| pBAD18 | Bacterial plasmid with Gentamycin resistance | Used to provide selectable resistance to *V. parahaemolyticus* prey strains in competition assays | Addgene |
| pBAD33 | Bacterial plasmid with chloramphenicol resistance | Used to provide selectable resistance to *V. natriegens* prey strains in competition assays | Addgene |
| pBAD33.1$^{F}$ | pBAD33.1 with a FLAG tag inserted at the 3' end of the MCS | Used for arabinose-inducible expression of proteins | (2) |

| | | | |
|---|---|---|---|
| pTfoX | $pBAD^{K}$/Myc-His plasmid containing the CDS of TfoX (VP1241) from *V. parahaemolyticus* RIMD 2210633 in frame with a C-terminal Myc-His tag | Used for the arabinose-inducible expression of TfoX | (4) |
| $pImmunity^{T2Rhs\text{-}NucRIMD}$ | $pBAD33.1^{F}$ containing the CDS of the $T2Rhs\text{-}Nuc^{RIMD}$ immunity protein (VP1518) from *V. parahaemolyticus* RIMD 2210633 in-frame with a C-terminal Flag tag | Used for the arabinose-inducible expression of the $T2Rhs\text{-}Nuc^{RIMD}$ immunity protein | This study |
| $pImmunity^{T2HydroRIMD}$ | $pBAD33.1^{F}$ plasmid containing the CDS of the $T2Hydro^{RIMD}$ immunity protein (VPA0348) from *V. parahaemolyticus* RIMD 2210633 in-frame with a C-terminal Flag tag | Used for the arabinose-inducible expression of the $T2Hydro^{RIMD}$ immunity protein | This study |
| $pT2Hydro^{RIMD/M65}$-i | $pBAD33.1^{F}$ plasmid containing the CDS of $T2Rhs\text{-}Nuc^{RIMD/M65}$ and its immunity protein (VPA0347 starting with methaionine 65 and VPA0348) from *V. parahaemolyticus* RIMD 2210633 in-frame with a C-terminal Flag tag | Used for the arabinose inducible expression of the $pT2Hydro^{RIMD/M65}$-i effector/immunity pair in BB22OP surrogate attacker strains in competition assays | This study |
| $pImmunity^{T2Rhs\text{-}NucBB22}$ | $pBAD33.1^{F}$ plasmid containing the CDS of the $T2Rhs\text{-}Nuc^{BB22}$ immunity protein (VPBB_RS07240) from *V. parahaemolyticus* BB22OP in-frame with a C-terminal Flag tag | Used for the arabinose-inducible expression of the $T2Rhs\text{-}Nuc^{BB22}$ immunity protein | This study |
| $pImmunity^{T2TmeBB22}$ | $pBAD33.1^{F}$ plasmid containing the CDS of the $T2Tme^{BB22}$ immunity protein (VPBB_RS25090) from *V. parahaemolyticus* | Used for the arabinose-inducible expression of the $T2Tme^{BB22}$ immunity protein | This study |

| | | | |
|---|---|---|---|
| | BB22OP in-frame with a C-terminal Flag tag | | |
| pImmunity$^{T2UnkwnBB22}$ | pBAD33.1$^{F}$ plasmid containing the CDS of the T2Unkwn$^{BB22}$ immunity protein (VPBB_RS18840) from *V. parahaemolyticus* BB22OP in-frame with a C-terminal Flag tag | Used for the arabinose-inducible expression of the T2Unkwn$^{BB22}$ immunity protein | This study |
| pImmunity$^{T2LipABB22}$ | pBAD33.1$^{F}$ plasmid containing the CDS of the T2LipA$^{BB22}$ immunity protein (VPBB_RS22625) from *V. parahaemolyticus* BB22OP in-frame with a C-terminal Flag tag | Used for the arabinose-inducible expression of the T2LipA$^{BB22}$ immunity protein | This study |
| pDM4 | a $Cm^{R}$ and $ori_{R6K}$-containing suicide vector | Used to generate deletions in *Vibrio* | (5) |
| pDM4:*hcp1*$^{RIMD}$ | pDM4 containing 1 kb downstream and 1 kb upstream of *vp1393* in its MCS | Used to delete *hcp1* in *V. parahaemolyticus* RIMD 2210633 | (3) |
| pDM4:*hcp2*$^{RIMD}$ | pDM4 containing 1 kb upstream and 1 kb downstream of *vpa1027* in its MCS | Used to delete *hcp2* in *V. parahaemolyticus* RIMD 2210633 | This study |
| pDM4:*hcp1*$^{BB22}$ | pDM4 containing 1 kb upstream and 1 kb downstream of *vpbb_rs06665* in its MCS | Used to delete *hcp1* in *V. parahaemolyticus* BB22OP | (2) |
| pDM4:*hcp2*$^{BB22}$ | pDM4 containing 1 kb upstream and 1 kb downstream of *vpbb_rs19920* in its MCS | Used to delete *hcp2* in *V. parahaemolyticus* BB22OP | (2) |
| pDM4:*T2Rhs-Nuc*$^{RIMD}$ | pDM4 containing 1 kb upstream and 1 kb downstream of *vp1517* in its MCS | Used to delete *T2Rhs-Nuc*$^{RIMD}$ in *V. parahaemolyticus* RIMD 2210633 | This study |
| pDM4:*T2Rhs-Nuc*$^{RIMD}$*-i* | pDM4 containing 1 kb upstream of *vp1517* and 1 kb downstream of *vp1518* in its MCS | Used to delete *T2Rhs-Nuc*$^{RIMD}$ and its downstream immunity gene in *V. parahaemolyticus* RIMD 2210633 | This study |

| | | | |
|---|---|---|---|
| pDM4:*T2Hydro*$^{RIMD}$ | pDM4 containing 1 kb upstream and 1 kb downstream of *vpa0347* in its MCS | Used to delete *T2Hydro*$^{RIMD}$ in *V. parahaemolyticus* RIMD 2210633 | This study |
| pDM4:*T2Hydro*$^{RIMD}$*-i* | pDM4 containing 1 kb upstream of *vpa0347* and 1 kb downstream of *vpa0348* in its MCS | Used to delete *T2Hydro*$^{RIMD}$ and its downstream immunity gene in *V. parahaemolyticus* RIMD 2210633 | This study |
| pDM4:*T2LipB*$^{RIMD}$ | pDM4 containing 1 kb upstream and 1 kb downstream of *vp0626* in its MCS | Used to delete *T2LipB*$^{RIMD}$ in *V. parahaemolyticus* RIMD 2210633 | This study |
| pDM4:*T2LipB*$^{RIMD}$*-i* | pDM4 containing 1 kb upstream of *vp0625* and 1 kb downstream of *vp0626* in its MCS | Used to delete *T2LipB*$^{RIMD}$ and its upstream gene in *V. parahaemolyticus* RIMD 2210633 | This study |
| pDM4:*T2Rhs-Nuc*$^{BB22}$ | pDM4 containing 1 kb upstream and 1 kb downstream of *vpbb_rs07235* in its MCS | Used to delete *T2Rhs-Nuc*$^{BB22}$ in *V. parahaemolyticus* BB22OP | This study |
| pDM4:*T2Rhs-Nuc*$^{BB22-}$*i* | pDM4 containing 1 kb upstream and 1 kb downstream of *vpbb_rs07235/40* in its MCS | Used to delete *T2Rhs-Nuc*$^{BB22}$ and its downstream immunity gene in *V. parahaemolyticus* BB22OP | This study |
| pDM4:*T2Tme*$^{BB22}$ | pDM4 containing 1 kb upstream and 1 kb downstream of *vpbb_rs07950* in its MCS | Used to delete *T2Tme*$^{BB22}$ in *V. parahaemolyticus* BB22OP | This study |
| pDM4:*T2Tme*$^{BB22}$*-i* | pDM4 containing 1 kb upstream of *vpbb_rs07950* and 1 kb downstream of *vpbb_rs25090* in its MCS | Used to delete *T2Tme*$^{BB22}$ and its downstream immunity gene in *V. parahaemolyticus* BB22OP | This study |
| pDM4:*T2Unkwn*$^{BB22}$ | pDM4 containing 1 kb upstream and 1 kb downstream of *vpbb_rs18835* in its MCS | Used to delete *T2Unkwn*$^{BB22}$ in *V. parahaemolyticus* BB22OP | This study |
| pDM4:*T2Unkwn*$^{BB22}$*-i* | pDM4 containing 1 kb upstream of *vpbb_rs18835* and 1 kb downstream of | Used to delete *T2Unkwn*$^{BB22}$ and its downstream immunity gene in *V.* | This study |

| | | | |
|---|---|---|---|
| | *vpbb_rs18840* in its MCS | *parahaemolyticus* BB22OP | |
| pDM4:*T2LipA*$^{BB22}$ | pDM4 containing 1 kb upstream and 1 kb downstream of *vpbb_rs22630* in its MCS | Used to delete *T2LipA*$^{BB22}$ in *V. parahaemolyticus* BB22OP | This study |
| pDM4:*T2LipA*$^{BB22}$*-i* | pDM4 containing 1 kb upstream of *vpbb_rs22625* and 1 kb downstream of *vpbb_rs22630* in its MCS | Used to delete *T2LipA*$^{BB22}$ and its upstream immunity gene in *V. parahaemolyticus* BB22OP | This study |
| pDM4:*T2LipB*$^{BB22}$ | pDM4 containing 1 kb upstream and 1 kb downstream of *vpbb_rs03020* in its MCS | Used to delete *T2LipB*$^{BB22}$ in *V. parahaemolyticus* BB22OP | This study |
| pDM4:*T2LipB*$^{BB22}$*-i* | pDM4 containing 1 kb upstream of *vpbb_rs03015* and 1 kb downstream of *vpbb_rs03020* in its MCS | Used to delete *T2LipB*$^{BB22}$ and its upstream gene in *V. parahaemolyticus* BB22OP | This study |
